# PyLabRobot: An Open-Source, Hardware Agnostic Interface for Liquid-Handling Robots and Accessories

**DOI:** 10.1101/2023.07.10.547733

**Authors:** Rick P. Wierenga, Stefan Golas, Wilson Ho, Connor Coley, Kevin M. Esvelt

## Abstract

Liquid handling robots are often limited by proprietary programming interfaces that are only compatible with a single type of robot and operating system, restricting method sharing and slowing development. Here we present PyLabRobot, an open-source, cross-platform Python interface capable of programming diverse liquid-handling robots, including Hamilton STARs, Tecan EVOs, and Opentron OT-2s. PyLabRobot provides a universal set of commands and representations for deck layout and labware, enabling the control of diverse accessory devices. The interface is extensible and can work with any robot that manipulates liquids within a Cartesian coordinate system. We validated the system through unit tests and several application demonstrations, including a browser-based simulator, a position calibration tool, and a path-teaching tool for complex movements. PyLabRobot provides a flexible, open, and collaborative programming environment for laboratory automation.

**Figure Abstract:** 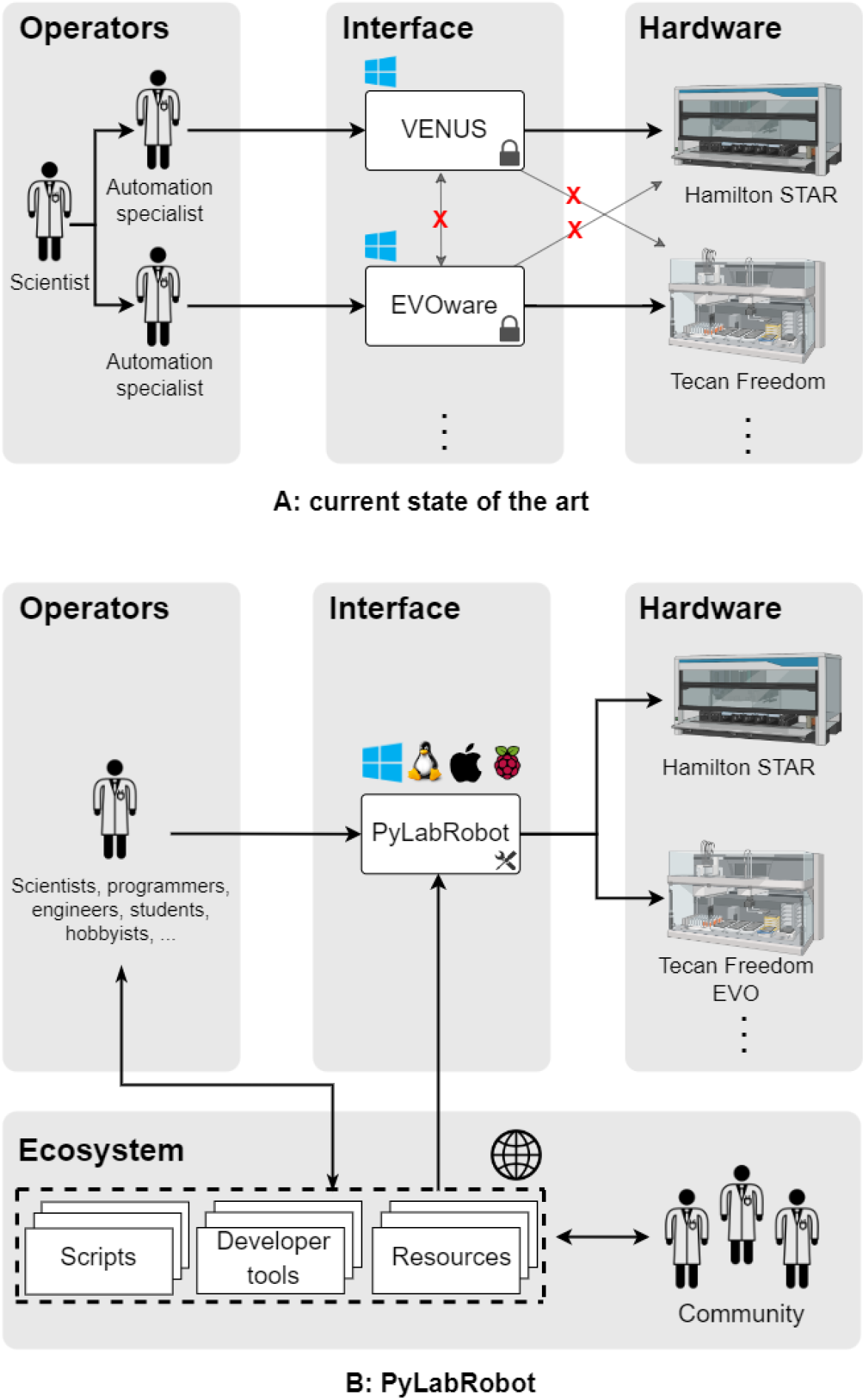

**PyLabRobot overcomes the limitations of proprietary robotic systems.** (a) Scientists with access to liquid-handling robots are currently limited by proprietary interfaces that require specialized knowledge, hinder cross-platform operability, and restrict sharing of methods among different robot types. For complex tasks, many researchers need assistance from a specialist familiar with their particular system, most notably when creating or editing protocols. (b) PyLabRobot (https://github.com/PyLabRobot/pylabrobot) offers a single interface that allows any person with basic Python skills to program diverse types of liquid-handling robots and share protocols freely, fostering a more collaborative and efficient research environment. The Python API makes it easy to interact with a large scientific computing ecosystem and allows users to leverage large language models for programming assistance.

## Introduction

Liquid-handling robots are versatile tools capable of automating a wide range of biology experiments. By replacing manual pipetting, they enable experiments to be executed in much higher throughput and with higher precision than is possible for humans. Robots are capable of performing many protocols, including mammalian tissue culture^1^, organoid culture^2^, hydrogel production^3^, directed evolution^4^, combinatorial drug screening^5^, plasmid assembly^6–8^, and high-throughput genomics sequencing^19^. Robotic platforms can also be integrated with computational resources such as machine learning models to perform closed-loop feedback control over iterated experiments^9–12^. However, current robots are limited by a ubiquitous reliance on proprietary software as the sole medium for creating and executing protocols, sharply constraining their flexibility, accessibility, and ability to integrate with external software resources.

In principle, the basic operations of loading disposable tips, aspirating and dispensing precise volumes of liquid, moving plates, and interfacing with accessories such as plate readers are highly similar across liquid-handling robots, suggesting that a protocol written for one robot could be readily adapted to another. In practice, idiosyncratic proprietary interfaces prevent developers from creating protocols that are interoperable across robots. Without easy sharing and adaptation, researchers are forced to write programs in proprietary formats using a limited library of tools that they cannot modify or extend. Proprietary graphical interfaces also prevent developers from freely integrating abstractions, tools, and libraries which are typical in other areas of software development.

Furthermore, because proprietary robot interfaces are typically not freely available to the public, there is little opportunity for newcomers to gain experience through educational, testing, or hobbyist usage. Each proprietary interface is an essentially unique and highly complex application that must be learned anew for each manufacturer, meaning proficiency does not readily translate across different robotic platforms. These factors sharply constrain the number of developers who can effectively program liquid-handling robots, resulting in an industry-wide personnel bottleneck for these roles.

Some of the problems of proprietary interfaces are mitigated by application programming interfaces (APIs) such as PyHamilton^13^ for Hamilton robots, the C# interface for Tecan Fluents, and the HTTP API for Opentrons. By decoupling the programming environment from the execution environment, APIs provide a useful layer of abstraction allowing for greater flexibility and the use of widely known programming languages. However, manufacturer-specific APIs still do not allow protocols and resources developed for one type of robot to be used on others. The issue of standardization to facilitate interoperability has yet to be adequately addressed by any existing interface. Several projects to create standardized lab automation interfaces have been created, such as SiLA^14^, LabOP^15^, Emerald Cloud Lab’s Symbolic Lab Language^16^, and Puppeteer^17^, but only the last of these supports command-level control of liquid-handling robots, but it is not publicly available or compatible with multiple types of robots.

Open-source applications outshine proprietary counterparts in their flexibility, accessibility, and compatibility with an extensive array of software development tools. Open source enables users to contribute changes that address their own context-specific needs and allows for integration with tools such as debugging and testing utilities^18^, and AI code assistants^19^. This allows for continuous improvement with features sourced from the widest possible pool of developers. These advantages have resulted in essentially all of the most widely used software development tools being open source, notably NumPy^20^, Jupyter Notebook^21^, PyTorch^22^, and countless others. Open-source software as such constitutes the foundation of modern software applications. An open-source interface for liquid-handling robots would dramatically improve protocol development efficiency and enable methods to be freely shared and reproduced across diverse operating system and hardware configurations. Here we present PyLabRobot, an open-source Python framework that provides a maximally flexible and accessible environment for programming liquid-handling robots and accessories.

## Results

PyLabRobot is designed to make developing and sharing protocols for liquid-handling robots and their accessories straightforward and accessible to users with a variety of skill levels. The platform is built to run on Windows, macOS, Linux and Raspberry Pi OS, allowing for accessibility to users of all major operating systems.

### Hardware Agnostic Interface

Users begin programming a laboratory device by instantiating a class from the interface layer, such as `LiquidHandler` or `PlateReader`, that is configured with a backend corresponding to their particular hardware platform. These expansive interfaces bridge the execution of commands, the deck layout, and the state of resources, ensuring that all specified locations exist and that requested operations will not cause conflict. Commands are passed to a backend in the execution layer responsible for translating them into instructions compatible with a specific device, and then conveying them to the machine through atomic, low-level commands in the form of firmware instructions or HTTP requests. Firmware instructions are communicated directly with machines using PyUSB^23^ and pylibftdi^24^ as platform-neutral wrappers and Python frontends for lower-level hardware libraries. All backends in the execution layer of a machine type inherit from a single abstract base class (ABC) to ensure uniformity. The open-source nature of PyLabRobot allows the entire community to incorporate a new instrument as soon as anyone writes and shares a backend for that specific piece of equipment.

By interfacing with equipment using atomic, low-level commands, PyLabRobot enables the use of interactive tools like REPLs (read-evaluate-print loops), IPython^25^, and Jupyter Notebooks^21^ to send individual commands to a robot to be executed in real time. This is in contrast to a typical protocol-based approach, in which robots can only be controlled by executing an entire script from beginning to end. An interactive interface has the advantage of enabling developers to have immediate feedback on the results of commands, allowing for rapid iteration and optimization of spatial positioning, liquid-handling parameters, and other physical variables.

### Deck Layout and Labware Model

Next, the user specifies their own unique configuration of labware on a robot deck with `Resource`, a class capable of capturing the attributes and methods defining most commonly used labware. Any configuration of labware on a robot deck can be represented as a set of instances of `Resource` and its subclasses, and can be modified at runtime. Each instance is defined by its `name` attribute, which serves as a unique identifier in the deck representation that every protocol can rely upon. These names typically refer to function rather than exact labware, which allows for robot agnostic protocols. For example, a protocol specifying aspiration from a resource named ‘bacteria lagoons’ will be interpreted by referencing the deck layout to obtain the specific labware definition.

Semantic relations between labware are stored in a directed, rooted tree (an arborescence), using the `children` and `parent` attributes of `Resource`, where the deck is the root of the tree (its `parent` is `None`). Spatial relations between resources are stored in the `location` attribute on `Resource`, which defines a resource’s location with respect to its immediate parent as a Cartesian coordinate. This allows locations to simultaneously be conceptualized as offsets. Absolute locations of resources are computed lazily by recursively adding a resource parent’s location, until the deck (root) is reached (Fig. 2). Key advantages of this approach include a straightforward implementation of moving resources (only the parent’s location has to be modified for all of its children to be likewise changed) and the ability to define resources (e.g., a plate with wells) without considering them in their broader context of a deck.

**Figure 1.**
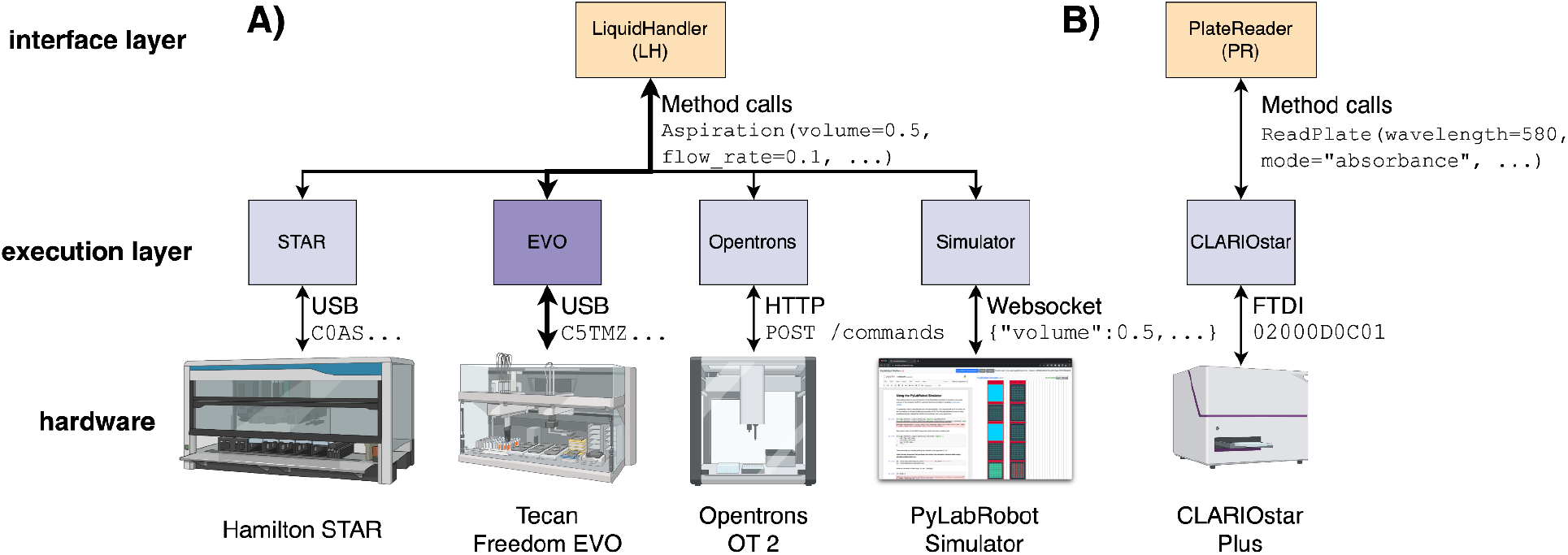
PyLabRobot software architecture. A) The operator always interacts with the same `LiquidHandler` interface layer. `LiquidHandler` communicates with the robot using a *backend* from the execution layer specific to the hardware in use. In this example, the Tecan Freedom EVO communication path is selected. Because the available hardware capabilities are conveyed to LiquidHandler by the execution layer, the user need only specify the configuration of the deck in order to translate methods developed for a different system to their own. B) Additional hardware devices mounted on the deck or nearby, such as plate readers, can be controlled using a similar architecture. Due to the open-source nature of PyLabRobot, the entire community can make use of a new hardware system as soon as one individual writes an appropriate backend.

**Figure 2.**
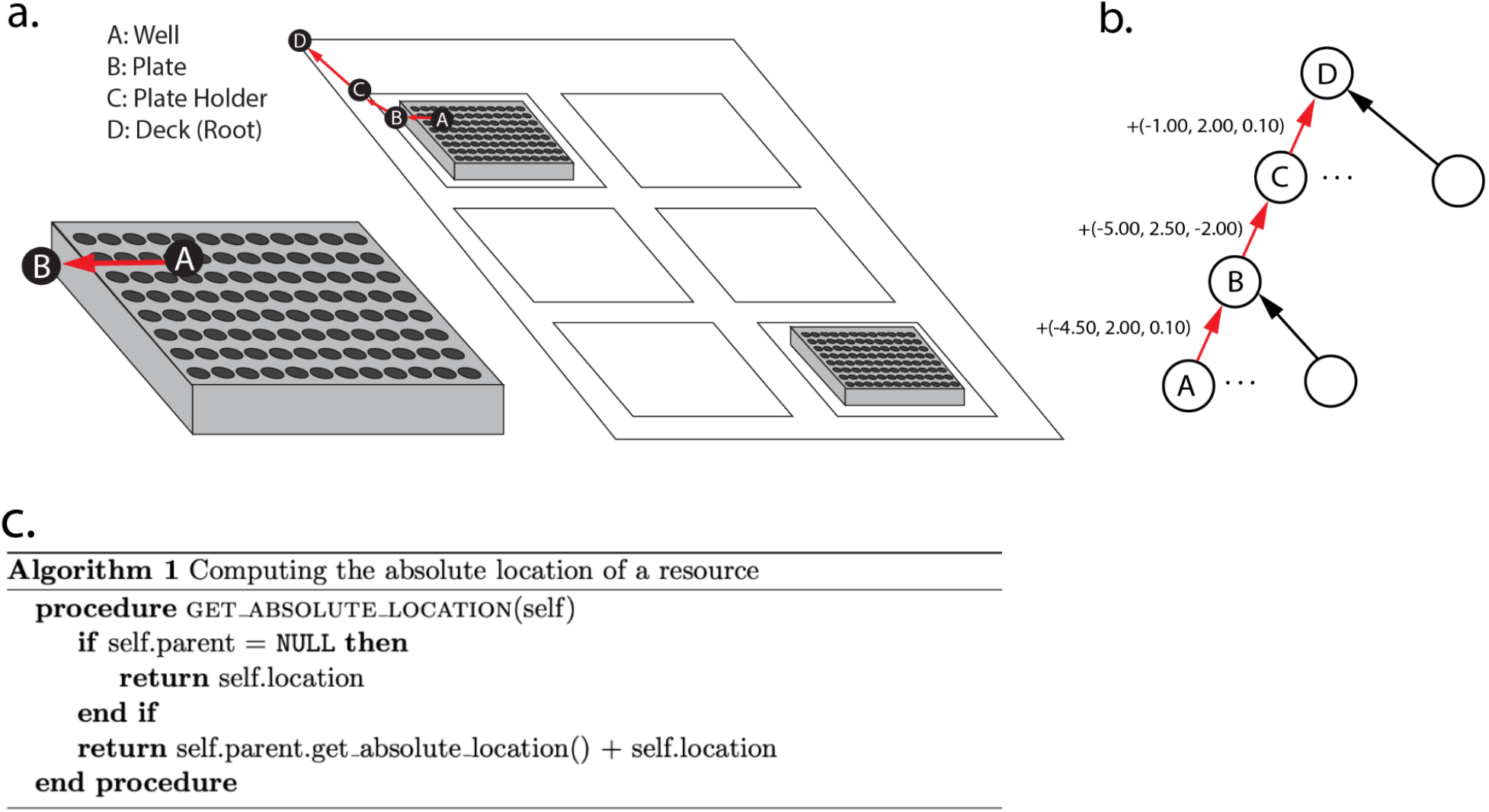
The Resource model tracking absolute position. (a) Visualization of the Resource model. The locations of wells are defined with respect to the bottom left of the plate. (b) The arborescence representing the computation of the location of resource A in resource D. The absolute location of the bottom left of the well is the sum of the location vectors between it and the deck. (c) Code identifying the location of a given instance of Resource. Users can define their own subclasses of Resource.

**Figure 3:**
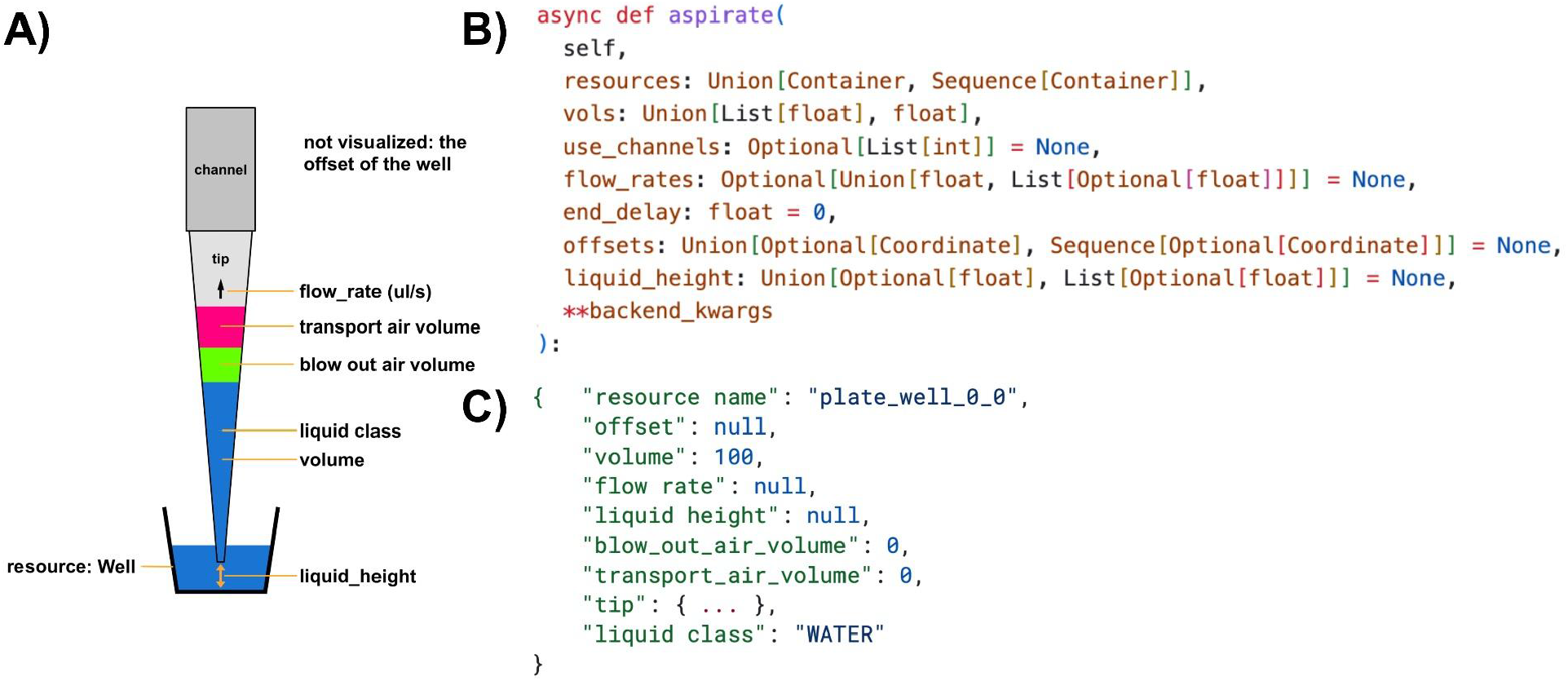
Basic liquid-handling operation and associated code. A) Schematic representation of an aspiration operation. The offset of the well is not pictured. B) The method header for aspiration on `LiquidHandler`. Type annotations help programmers use the library. Optional values need not be specified and the best values may be chosen by a backend. C) Standard form operations can be serialized into and deserialized from JSON format.

Subclasses of `Resource` capture additional behavior and constraints of particular labware types. The following are the most important subclasses of `Resource` in PyLabRobot:

- `Deck` serves as the root of all resources in the deck layout arborescence described above. Exploiting the similarity between resource-to-resource relations, the `Deck` class is a subclass of `Resource` with a few minor modifications. First, it maintains a mapping of a resource’s name to a reference to that resource. This allows random access of resources in *O(1)*, which is particularly useful when checking for naming collisions. The choice to only include this functionality in this subclass was made for space efficiency. Second, the deck calls a callback method on assignment of resources, so that LH can listen for changes in deck geometry and, if necessary, communicate those to a backend. Each robot implementation currently includes a subclass of `Deck` to define its deck geometry. The `HamiltonDeck` subclass additionally includes a method to load VENUS’ lay files into the PyLabRobot model.
- `Carrier` is an abstract base class for tip rack and plate carriers. Both provide a fixed number of well-defined spots, in which at most 1 tip rack or plate can be placed respectively. `Carrier` mirrors this constraint by overriding the `assign_child_resource` method to require a fixed spot, and raising an error if the requested spot is already occupied.
- `Container` is an ABC for resources that contain liquid, and uses a volume tracker to keep track of the used and free volume. Subclasses include `Trough` and `Well`. Volume trackers will be explained in more detail later on.
- `ItemizedResource` is a generic ABC for resources that contain children in a uniform grid configuration, most notably `Plate` and `TipRack`. It provides convenience methods for indexing these children using conventional alphanumeric notation (<row letter><column number>, *eg “A1”*) and integer indices, as well as a method for traversing them. `Plate` and `TipRack` have an associated type of `Well` and `TipSpot` respectively.

The user can subclass `Resource` as well as the abovementioned subclasses to further encapsulate specific labware properties. For example, a user may subclass `ItemizedResource` with an associated subclass `Container` to represent a rack of vials.

Each instance of `Resource` can serialize and deserialize itself and its children to and from JSON format using the `serialize` and `deserialize` methods respectively. The recursive and flexible implementation means third party developers rarely have to write their own deserializer. Thanks to JSON’s flexibility and versatility, resource definitions and labware configurations can be easily shared with a wide range of external software.

### Standardized operations

All liquid-handling operations in PyLabRobot are implemented using four universal and foundational commands: aspirate, dispense, tip pickup, and tip drop. These commands correspond to methods on `LiquidHandler` and `LiquidHandlerBackend` (`pick_up_tips`, `drop_tips`, `aspirate`, `dispense`), as well as four dataclasses to transfer the data between them (`Pickup`, `Drop`, `Aspiration`, `Dispense`). Multiple instances of such a class may be passed to a backend method in a list, with the expectation that they are executed in a single pass. If many operations are carried out simultaneously on a single tip rack or plate using a special pipetting head, such as the CoRe-96 head on Hamilton’s STAR, the *-TipRack* and -*Plate* variants of each of the methods and classes should be used (`PickupTipRack`, `DropTipRack`, `AspirationPlate`, `DispensePlate`). These variants differ in how they store tips as a list rather than a single element, for example. In addition, a `Move` operation exists to capture the parameters of resource movement. Some parameters need not be specified explicitly (i.e. they may be `None`), in which case a backend may choose the most appropriate values.

Composition of the unit operations, such as for example the discard operation (drop tip in trash) and transfer operations (combined aspirate and dispense), is done at the `LiquidHandler` level and above. The responsibilities of each backend are minimized to make adding new robot models as easy as possible.

### Monitoring and parallelization

PyLabRobot uses “Tracker” objects to track and validate the physical state of the robot, including the presence of tips in a tip rack and liquids in wells and other liquid containers. This information is used to catch potential errors before executing operations on hardware, which can be costly, as well as to facilitate higher level features like `return_tips`. Trackers use a transaction pattern when validating and saving operations. Before a command is sent to a machine, operations are validated against the currently queued state and present state. If validation passes, the operation is added to the queue. Many operations may be queued in this manner. After that, the operations in the queue are executed in order. If all operations execute successfully on the machine, the entire queue is either committed (the current state is updated to be the queued state), otherwise changes are rolled back (the queued actions are deleted).

Tip trackers track the presence of tips on the pipetting head and tip spots in a tip rack, and can distinguish between fixed or disposable tips; volume trackers track the liquids in liquid containers such as wells and mounted tips.

Executing steps of automation protocols typically takes a meaningful amount of time, during which other tasks can or must be executed^4,26^. Using Python’s **asyncio** library makes it easy to compose protocols that run steps in parallel (Fig. 4).

**Figure 4:**
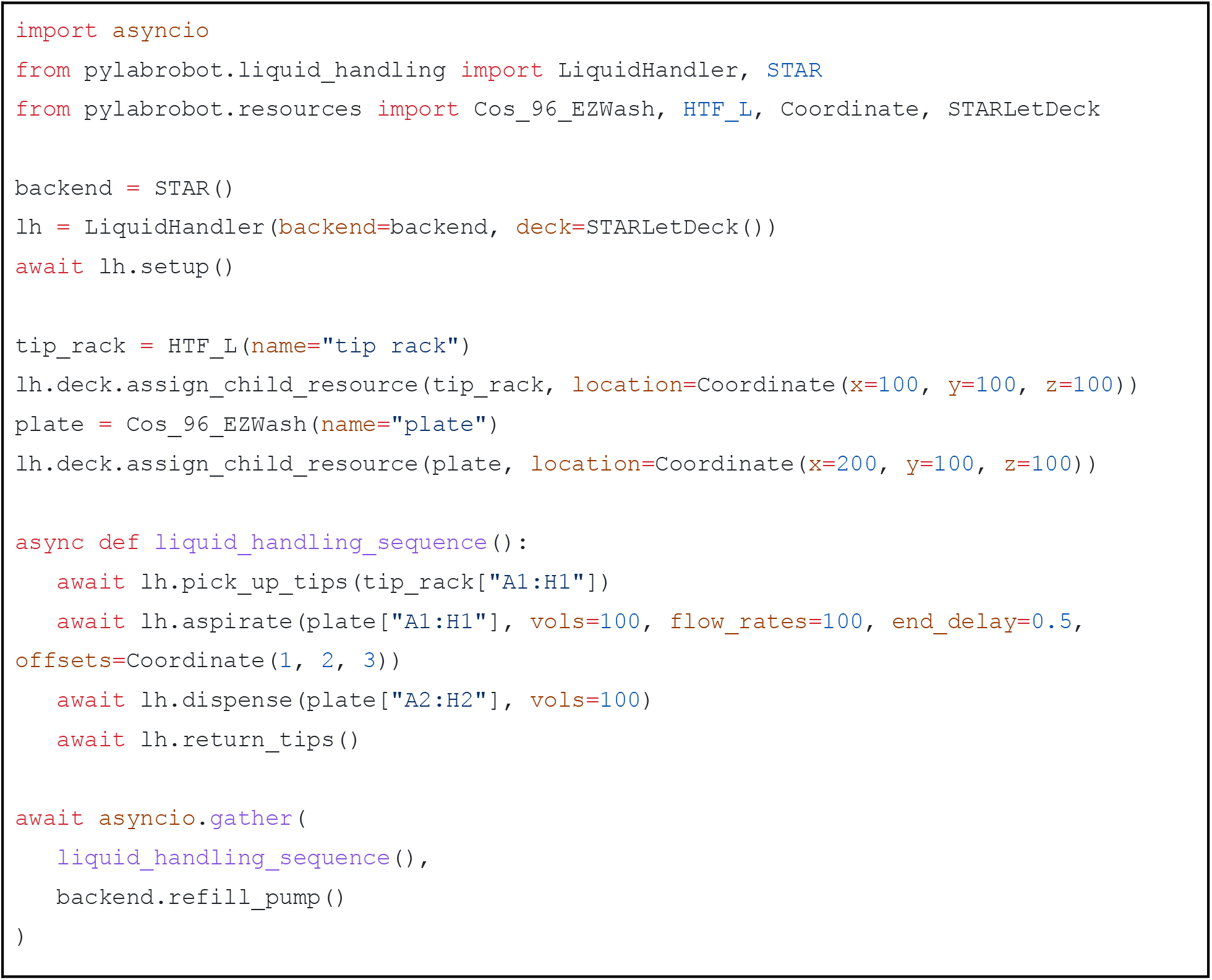
PyLabRobot example script. This will move 100uL of liquid from the first to second column. The tips are returned to their pickup location. The refilling of the washer station, a backend-specific activity, is done simultaneously. The program ends after the longest running task finishes.

**Figure 5.**
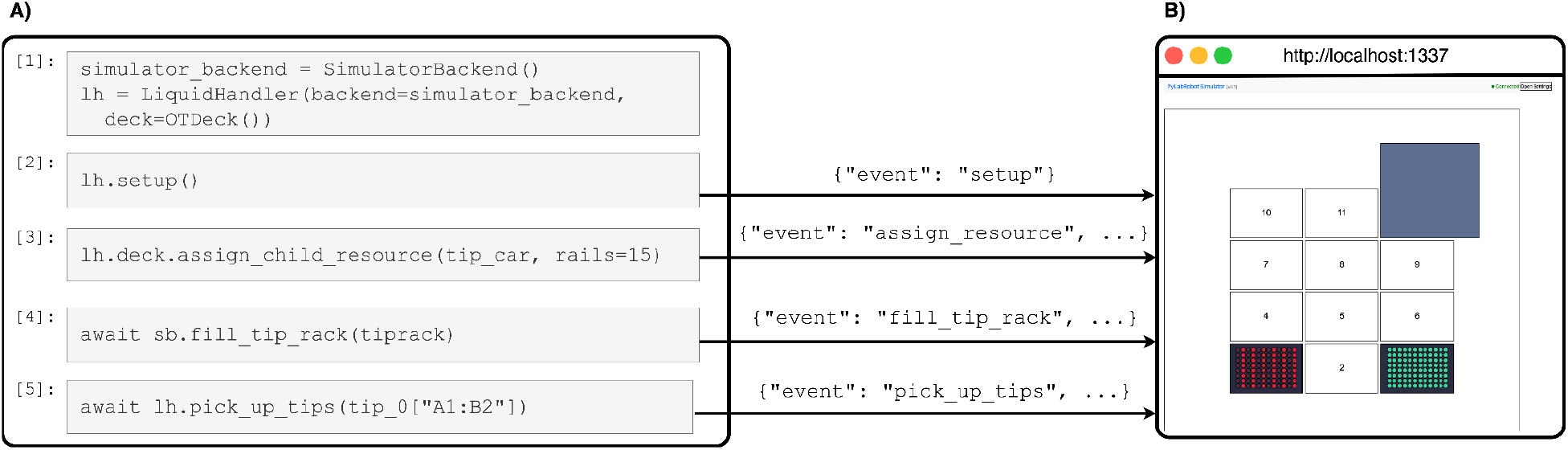
The simulator. (a) A Jupyter notebook that interactively uses PyLabRobot. (b) The simulator running in a web browser to visualize the resources during a run. Tip presence and volumes are visualized in the simulator. Parts of commands sent over a websocket connection during execution of each cell are visualized; command responses are not visualized.

### Simulator

To lower the barrier of entering the field of lab automation, as well as to more easily validate the correctness of the library and methods, a browser based simulator was created to allow people to use PyLabRobot without having to have any specialized hardware. This simulator is part of the open source package and can be run locally. Alternatively, a hosted version of the simulator is available at simulator.pylabrobot.org.

The simulator is controlled using the `SimulatorBackend`, a subclass of `WebsocketBackend` with additional support for simulator specific functionality. For example, this backend provides methods for updating the “physical” state of the deck, like placing tips and liquids on the deck.

The browser based part of the simulator mirrors the robot agnostic resource model, and is dynamically constructed and modified during a simulator run. This, combined with the fact that the backend passes along all information received by LiquidHandlerBackends, means the simulator works with any robot for which a deck model exists in PyLabRobot. The only code a developer has to write to add a new robot to the simulator is to enforce machine specific constraints, such as the specific configuration of pipetting channels.

### Graphical labware layout editor

A graphical labware layout editor was developed to make designing labware layouts faster and more accessible. The labware editor reuses the same UI components used in the simulator for maintainability and visual consistency. The editor can also be used to edit the initial state of a deck by editing the presence of tips in tip racks and liquids in wells and other liquid containers. This data is saved to a layout and state JSON file on disk after validation, both of which can be loaded into PyLabRobot.

### Applications

#### Demos

The PyLabRobot Art Studio (https://github.com/rickwierenga/pylabrobot-art-studio), an application that prints users’ 12 by 8 drawings using watercolor paint, demonstrates PyLabRobot’s ability to execute complicated liquid handling operations on an Opentrons. The demo includes a web server and custom UI. The demo reuses tips when possible, which can be thought of as a way to eliminate contamination. Both the integration with external libraries and the dynamic pattern of liquid handling would be (virtually) impossible to achieve with traditional software.

The Game of Life demo (https://github.com/rickwierenga/plr-game-of-life) demonstrates the plate reader integration in PyLabRobot by running a Game of Life simulation^27^, using wells as the exclusive memory medium by using crystal violet dye to indicate “living wells”.

#### Labware Position Calibration, Path Teaching, and Interactivity

A common challenge in creating custom deck layouts is calibrating positions of labware to a set of coordinates in 3D space. To alleviate this, we created a tool called PyLabRobot Resource Locator Program (PLR-RLP, https://github.com/PyLabRobot/resource-locator-program) that exploits PyLabRobot’s interactive control over a machine to move a pipetting tip to an arbitrary location and assign not-yet-located labware at that location. In addition, this same program can be used to pick up and release resources at arbitrary locations, and to design the intermediate traversal path. The program can be controlled using the keyboard, a GUI, and a video game controller.

#### LLM Assistant

Programmatic robot interfaces are naturally conducive to integration with large language model (LLM) assistants. One common use of LLM coding assistants is to provide a natural language prompt for which the assistant writes code. This can enable users who are less skilled with programming, or even completely unskilled with Python, to work with liquid-handling robots^28^. To provide a wider group of biologists with access to laboratory automation, OpenAI’s GPT-3.5 model was provided with examples of PyLabRobot code, and then used to convert natural language prompts directly into usable robot code. **Table 3** provides a number of prompts and outputs from OpenAI GPT-3.5 model after being given examples of PyLabRobot code. A Jupyter notebook containing the example data can be found at: https://github.com/stefangolas/PyLabRobot_LLM_Example.

**Table 1.** Package organization.

| Subpackage Name | Description |
| --- | --- |
| **liquid_handling** | Base classes and implementations for communicating with liquid-handling robots and tracking robot state |
| **liquid_handling.backends** | Backends for translating liquid-handling commands to robot-specific commands |
| **liquid_handling.liquid_classes** | Classes that specify default parameters when handling different types of liquids |
| **plate_reading** | Tools for working with plate readers |
| **resources** | Base classes and implementations for specifying deck and labware attributes and methods |
| **server** | HTTP server that receives and executes liquid handling commands |
| **utils** | Generic high-level utilities |

**Table 2.** Basic liquid-handling operations available through PyLabRobot. Composite operations exist in the interface layer for reusability, but are implemented on the backend as multiple fundamental operations to minimize the responsibility in backends, making adding new backends and devices easier.

| LH method | Description | Backend method(s) |
| --- | --- | --- |
| `pick_up_tips` | Pick up tips from a tip rack | `pick_up_tips` |
| `drop_tips` | Put tips back in a tip rack, reuse allowed | `drop_tips` |
| `discard_tips` | Dispose of the tips, reuse not allowed | `drop_tips` |
| `return_tips` | Return the mounted tips to where they were picked up | `drop_tips` |
| `aspirate` | Aspirate from a container | `aspirate` |
| `dispense` | Dispense to a container | `dispense` |
| `transfer` | Transfer liquid from one container to another | `aspirate, dispense` |
| `pick_up_tips96` | Pick up 96 tips from a tip rack using a single head | `pick_up_tips96` |
| `drop_tips96` | Drop 96 tips to a tip rack with a single head, reuse allowed | `drop_tips96` |
| `discard_tips96` | Dispose of 96 tips to the trash location, reuse disallowed | `drop_tips96` |
| `return_tips96` | Return all 96 tips to the tip rack where they were picked up | `drop_tips96` |
| `aspirate96` | Aspirate from a container using 96 channels at once | `aspirate96` |
| `dispense96` | Dispense to a container using 96 channels at once | `dispense96` |
| `move_resource` | Move a resource to a new location | `move_resource` |
| `move_plate` | Move a plate to a new location | `move_resource` |
| `move_lid` | Move a plate lid to a new location | `move_resource` |

**Table 3.** Examples of natural language prompts to control PyLabRobot. GPT-3.5 can generate PyLabRobot code to perform desired tasks. The modular structure and defined deck layouts of PyLabRobot improve reliability, while the simulator permits testing prior to robot use.

| Prompt | Code Output |
| --- | --- |
| aspirate 20uL from the first and last wells in column 2 | `asp_poss = [plate[8], plate[15]] lh.aspirate([asp_poss], vols=[20]*2)` |
| aspirate 100uL from the last four wells of column 4 | `asp_poss = [plate[i] for i in range(28,32)] lh.aspirate([asp_poss], vols=[100]*4)` |
| pick up tips and aspirate 200uL from column 1 and dispense to column 3 | `tips_poss = [tip_0[i] for i in range(8)] lh.pick_up_tips(tips_poss) asp_poss = [plate[i] for i in range(8,16)] lh.aspirate(asp_poss, vols = [200]*8) disp_poss = [plate[i] for i in range(16,24)] lh.dispense(disp_poss, [200]*8)` |
| pick up tips and aspirate 200uL from column 1 and dispense 50uL to columns 5,6,7 | `tips_poss = [tip_0[i] for i in range(8)] lh.pick_up_tips(tips_poss) asp_poss = [plate[i] for i in range(8)] lh.aspirate(asp_poss, vols = [200]*8) disp_cols = [4,5,6] for col in disp_cols: disp_poss = [plate[idx] for idx in range(8*col,8*col+8)] lh.dispense(disp_poss, [50]*8)` |

## Discussion

As an open-source library capable of operating diverse liquid-handling robots and other laboratory hardware, PyLabRobot provides the laboratory automation community with a unique opportunity to share protocols and benefit from one another’s work, improving efficiency and access. By allowing researchers to write protocols using Python, one of the most widely accessible programming languages, the library simultaneously lowers the barrier to entry for researchers who have less experience in writing software and allows experts to switch to a new hardware platform without learning a new interface and rewriting all of their methods. Python is distinctly advantageous for this context because it has an extremely large ecosystem of libraries for use-cases that synergize with lab automation, particularly data analytics, bioinformatics and machine learning^29^ .

The flexibility of an open-source framework allows for easy integration of libraries, tools, and abstractions into developer workflows. Encapsulation of code into reusable abstractions is a highly productive pattern that is ubiquitous in software development but has only been minimally deployed within lab automation. Code reusability has benefits for developer productivity and experimental scale and reproducibility, and the open-source nature of PyLabRobot enables the nucleation of a developer ecosystem that facilitates maximum distribution of shared resources.

The advent of large language models (LLMs) solidifies the benefits of a universal programmatic interface to liquid-handling robots. These models are capable of translating natural language prompts into Python code, allowing experimentalists with negligible programming experience to automate protocols. LLM agents can design and execute experiments on robots by parsing scientific literature. The modular structure of PyLabRobot enables such an agent to autonomously design and execute experiments on a range of liquid-handling robots with maximum flexibility and interoperability.

Already compatible with Hamilton, Tecan, and Opentrons liquid handling robots and BMG plate readers, PyLabRobot can readily be extended to interface with other liquid-handling robots (including open-source models such as the EvoBot^30^) and lab automation equipment, enabling them to be run from any operating system and interact with a huge computing ecosystem. As additional hardware and protocols are added to the repertoire, labs will obtain greater benefit from using PyLabRobot, incentivizing manufacturers to provide PyLabRobot-compatible drivers for their equipment. One day, we hope all laboratory automation equipment will be operable using a highly flexible and OS-agnostic interface, enabling all users to benefit from a massive library of shared protocols that can readily be adapted for any purpose.

## Methods

### Software development best practices

The PyLabRobot project, as a foundation for higher level libraries and applications, must be stable. To ensure quality across the board, several software development best practices are used. The library makes use of automated systems, running on GitHub Actions and developer’s machines. Releases of the library are automatically pushed to the Python Package Index (PyPI) so that they may easily be installed by the user.

#### Static code analysis

Mypy^31^ is used to enforce static typing rules, as a way to mitigate some of the downsides of Python’s dynamic typing system as well as provide the user with additional usage information. PyLint^32^ is used to statically analyze code for errors and bad practices, and to ensure consistent code style, which is particularly helpful as the community of contributors to the project grows.

#### Automated testing

Each commit must pass a battery of tests before it is merged into the central repository. Tests are executed using Pytest^18^. At the execution layer, unit tests are used to ensure that user input produces consistent machine level commands, and, if possible, that they are transmitted correctly (after sending the commands they are no longer PyLabRobot’s responsibility). This means all tests can be run without requiring access to any specialized hardware.

#### Documentation

Documentation (available at docs.pylabrobot.org) is generated using Sphinx^33^, one of the most popular tools to generate documentation from Python docstrings. Pylint is set up to force each public class and all methods and functions to have a docstring, thereby making sure the entire library is well documented. This has the additional benefit of keeping code and documentation closely related. The autodoc extension is used to automatically and recursively generate documentation for various modules and submodules. Using MyST-NB^34^, integrated tutorials can be written in an interactive format (Jupyter Notebook^21^), allowing the end user to easily run the provided examples in their own laboratory.

#### Code simulation

Traditionally, all development for liquid handling robots required ownership of a physical machine, which hindered learners and development tremendously. It is still true that some facets of development will require physical access to a robot. This will be mitigated by placing trust on authorities such as manufacturers and through peer review. The simulator is a powerful tool that aids this process.

#### Abstract base classes

PyLabRobot provides a number of classes that define the general behavior of concepts such as backend, robot deck, or deck resource through abstract base classes (ABCs). Classes that define specific implementations of one of these abstractions are defined as subclasses which inherit from the corresponding ABC. Defining classes to represent abstractions of each concept enables functionality to be generalized and enforced across each of the particular subclasses. This enforces standardization in how diverse instruments and systems are handled, so that developers can expect predictable behavior from PyLabRobot resources (such as robot backends), even for future integrations. The standard may be updated to accommodate future developments in lab automation.

### Community

The greatest benefits of open-source software require a flourishing community to share code and assist one another with problem-solving and troubleshooting. We consequently built and now host an active forum to promote knowledge sharing in lab automation (forums.pylabrobot.org). Described by one user as a “huge improvement over the popular private channels of communication”, it has grown to feature 20+ posts per day on a wide variety of laboratory automation topics.

### Labware Libraries

Labware definitions from Hamilton, Tecan, and Opentrons software have been translated into the hardware-agnostic labware model described in Results. Hardware agnosticity has been verified by using Hamilton resources on an Opentrons robot and vice versa. This library of hardware agnostic labware definitions could serve as the basis for a hosted universal labware database that would facilitate interoperability across robot platforms.

### Web connectivity

Web based protocols provide a powerful, versatile and extendible interoperability layer for communicating between different programs and computers. Included in PyLabRobot is an application level protocol for interfacing with liquid handling robots and plate readers, as well as a client and server implementation. The `SerializingBackend` is available as a transmission-layer neutral encoder. The two web protocols supported by PyLabRobot are HTTP and Websockets^35^, for which transmission is implemented by the `WebsocketBackend` and `HTTPBackendServer` backends. An HTTP front end is provided which decodes the data and sends it to an instance of `LiquidHandler`. It must be explicit that this protocol can be implemented by any program, meaning PyLabRobot may run just on the server or client side, on both, or nowhere.

### STAR Backend

The `STAR` backend provides an operating system agnostic interface to the Hamilton STAR and STARLet robots that has been verified to work with Windows, macOS and Raspberry Pi OS. The backend communicates directly with the robot’s firmware over a USB connection. STAR backend firmware commands were based on those used by the Venus application to communicate with the STAR, and are documented in Hamilton reference manuals. The PyLabRobot STAR interface works with multiple USB drivers thanks to PyUSB^21^. During testing, the libusbK driver was used on Windows, libUSB on macOS and the driver provided by the operating system on Raspberry Pi OS (Debian Linux). This creates a communication channel to the robot that is completely open source.

### EVO Backend

The `EVO` backend provides an operating system agnostic interface to the Tecan Freedom EVO and has been validated to work on Windows and macOS. Similar to `STAR`, `EVO` uses firmware command strings based on those used by the EVOWare application and documented in the manufacturer’s reference manuals.

### Opentrons Backend

The Opentrons backend uses the Opentrons HTTP API, for which a Python wrapper was written (https://github.com/rickwierenga/opentrons-python-api). The server runs on the onboard Raspberry Pi, and PyLabRobot can either run on the onboard computer or an external computer. This provides an advantage over the usual mode of operation where entire protocols are stored on device, potentially making having a single source of truth the user’s concern, limiting the ecosystem a protocol may interact with, and typically forcing the user to run entire protocols at once rather than interactively.

Since the Opentrons API currently has to maintain its own deck model, PyLabRobot’s deck model is mirrored in the background using the `assigned_resource_callback` and `unassigned_resource_callback` callback methods.

## Acknowledgements

The authors would like to thank Dana Gretton, Emma Chory, Jon Bloom, Alvaro Cuevas, Eric Sindelar, Ben Ray, Priyanka Raghavan, and Wenhao Gao for constructive feedback. We would also like to thank Ben Gregor, Eyal Perry, David Kong, Priyanka Raghavan, Wenhao Gao, and Christian Ulmer for assistance with procuring documentation and hardware vital to the success of this project. We are deeply grateful for support from Reid Hoffman, the Open Philanthropy Project, the National Institutes of Health (grants no. R21AI158169 and DP2AI136597), and the MIT Media Lab.

## Author contributions

R.W. and S.G. conceived of the idea. R.W. and W.H. developed the PyLabRobot software library with advice from S.G. and K.M.E. R.W. and S.G. validated the software and developed the demonstrations. R.W., S.G., C.C. and K.M.E. procured the devices and materials. R.W., S.G. and K.M.E. wrote the paper with input from all authors. K.M.E. procured funding.

